# *Theobroma cacao* genome sequence reveals insights into flavonoid biosynthesis

**DOI:** 10.1101/2024.11.23.624982

**Authors:** Maria Fernanda Marin Recinos, Sarah Winnier, Kiersten Lagerhausen, Blessing Ajayi, Katharina Wolff, Ronja Friedhoff, Chiara Marie Dassow, Nancy Choudhary, Julie Anne Vieira Salgado de Oliveira, Boas Pucker

## Abstract

*Theobroma cacao* is well known for its role in producing cacao. Many different cacao varieties are cultivated in equatorial regions, each distinguished by unique phenotypic traits. One of these traits is pod color variation, which has been linked to differences in anthocyanin content. Anthocyanins are produced by a branch of the flavonoid biosynthesis pathway. In this study, two high-contiguity *T. cacao* genomes, BRAU1 and BONN1, were sequenced and assembled and used to investigate genes and structural variants associated with intraspecific variation in flavonoid biosynthesis. Comparative analysis identified structural variations affecting six candidate flavonoid biosynthesis genes, including a 1026 bp deletion at one of the *DFR* loci in BRAU1, BONN1, and Matina accessions.

## Introduction

*Theobroma cacao*, a diploid species (2n = 2x = 20) containing ten chromosomes, is thought to be native to South American rainforests and part of Central America. It is a tropical perennial tree belonging to the *Malvaceae* family and is globally recognized for the commercial value of its beans, primarily used in the production of chocolate and other derivatives (Motamayor *et al*. 2002). The high demand for cocoa has made *T. cacao* one of the most important commercial crops in the world (Voora *et al*. 2020; FAO 2022), and it is now cultivated in equatorial regions and zones with tropical climate (Cadby & Araki 2021).

Historically, cacao varieties have been classified into three groups or subspecies: Criollo, Forastero, and Trinitario (Cheesman 1944). More recent research has proposed a division into ten groups, namely Amelonado, Contamana, Criollo, Curaray, Guianna, Iquitos, Marañon, Nacional, Nanay, and Purús (Nousias *et al*. 2024). Criollo has been the most extensively studied due to its high degree of homozygosity (Motamayor *et al*. 2002). The Criollo group is primarily found in the Americas and is known for producing red fruits, smooth skin, and low seed yield (Nieves-Orduña *et al*. 2023). This variety is commercially valued for its high-quality beans, which are described as having a delicate floral or fruity taste with low acidity and bitterness. However, Criollo trees generally present lower yields and higher susceptibility to diseases, while the Forastero variety is more vigorous, with higher seed productivity but a stronger bitter taste (Toxopeus 2001; Gopaulchan *et al*. 2019). Forastero trees are known to produce fruits with a rough surface and a coloration ranging from yellow to red. This variety is native to the Amazon basin and primarily cultivated in regions of central and western Africa (Toxopeus 2001). Moreover, the Trinitario variety, a hybrid of Criollo and Forastero, combines desirable traits from both parent varieties, offering a balance of flavor and resilience (Toxopeus 2001).

Despite the traditional classification based on morphology and geographical origin, recent studies have shown no clear genetic basis for these distinctions (Motamayor *et al*. 2002). Some Forastero individuals are as genetically distant from each other as from Criollo varieties (Motamayor *et al*. 2002), highlighting the complexity of cacao genetics. Due to the extensive range where cacao is now grown, the cacao genetic pool has diversified. Considerable efforts have been made to understand the genetic diversity behind specific geographical groups have been made, revealing how diverse the cacao gene pool is and the power of gene flow when introducing a different variety (Motamayor *et al*. 2008; Gopaulchan *et al*. 2019, 2020; González-Orozco *et al*. 2022). This has made accurate taxonomic classification of *T. cacao* varieties and subvarieties challenging for population geneticists and botanists.

Beyond its significance in the food industry, *T. cacao* has been studied for its potential medicinal properties. Historical records, such as the Florentine Codex in 1590, document the use of cacao in treating fever and heart conditions (Rusconi & Conti 2010). Since 1996 alone, there have been over 38 studies of the cardiovascular benefits of cacao in humans, with many attributing these effects to its high polyphenol content, which acts as an antioxidant when ingested (Rusconi & Conti 2010). *T. cacao* beans are known to be a rich source of polyphenols, making up about 10% of the dry weight of the bean (Rusconi & Conti 2010). The largest and most diverse group of phenols present in *T. cacao* beans is flavonoids (Oracz *et al*. 2015). There are several known groups of flavonoids found within *T. cacao* beans including catechins, epicatechins, and anthocyanins (Acosta-Otálvaro *et al*. 2022). These compounds are specialized metabolites that are stored within the vacuoles primarily in cotyledons (Oracz *et al*. 2015).

The flavonoid biosynthesis pathway has been studied in detail in model plant species like *Arabidopsis thaliana* (Winkel-Shirley 2001; Grotewold 2006; Appelhagen *et al*. 2014). The biosynthesis of flavonoids (**Figure 1**) begins with the general phenylpropanoid pathway, followed by the formation of chalcone through the combination of 3 malonyl-CoA molecules and 4-coumaroyl-CoA, catalyzed by the chalcone synthase (CHS) (Ferrer *et al*. 1999). Next is the isomerization of chalcone to naringenin by the chalcone isomerase (CHI) (Jez *et al*. 2000). The pathway’s complexity is caused by many branching reactions, resulting in the formation of various flavonoid subclasses, including flavones, flavonols, anthocyanins, and proanthocyanidins (Winkel-Shirley 2001; Grotewold 2006). The anthocyanin and proanthocyanidin biosynthesis form branches of the flavonoid biosynthesis that compete with the formation of flavonols (Winkel-Shirley 2001; Grünig *et al*. 2025). The flavonol biosynthesis pathway branches off by the conversion of dihydroflavonols to flavonols, catalyzed by the flavonol synthase (FLS) (Owens *et al*. 2008). Flavonols are then decorated with sugar moieties catalyzed by glycosyltransferases (Offen *et al*. 2006; Yonekura-Sakakibara *et al*. 2014; Ishihara *et al*. 2016) and believed to be stored in the central vacuole. Flavonols are often reported as UV protective compounds but also appear to have regulatory activities (Williams *et al*. 2004; Pollastri & Tattini 2011; Naik *et al*. 2024).

**Figure 1.**
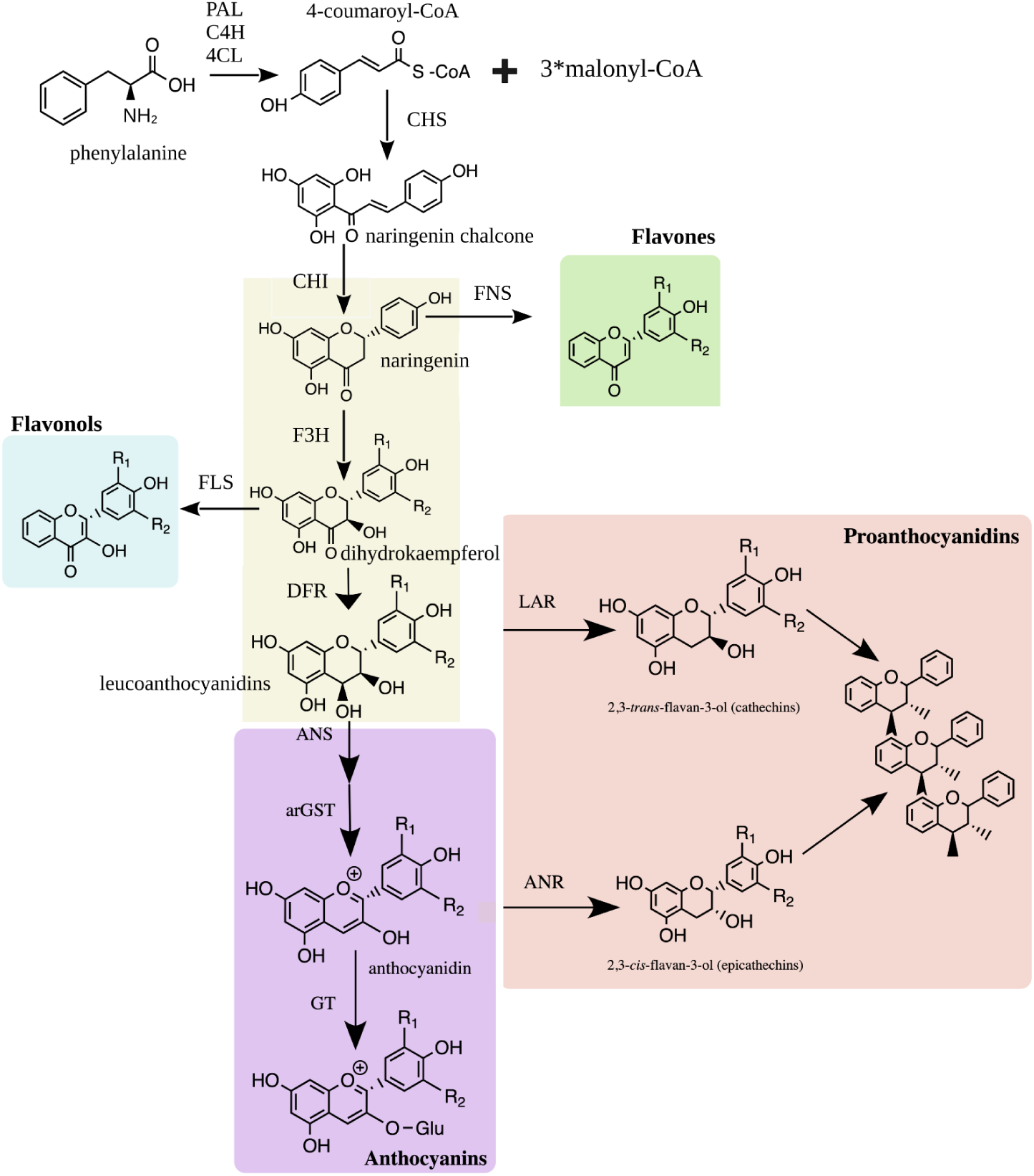
Schematic representation of the general flavonoid biosynthesis pathway. Enzyme names are abbreviated as follows: PAL - phenylalanine ammonia-lyase, C4H - cinnamic acid 4-hydroxylase, 4CL − 4-coumarate-CoA ligase, CHS - chalcone synthase, CHI - chalcone isomerase, FNS - flavone synthase, F3H - flavanone 3-hydroxylase, FLS - flavonol synthase, DFR - dihydroflavonol 4-reductase, ANS - anthocyanidin synthase, arGST - anthocyanin-related glutathione S-transferase, GT-dependent anthocyanidin 3-O-glycosyltransferase, LAR - leucoanthocyanidin reductase, and ANR - anthocyanidin reductase.

Anthocyanins are well known as pigments providing colorful hues to plant tissues, but also play roles in protection against UV light, reactive oxygen stress, and many other processes (Grünig *et al*. 2025). In *T. cacao*, variations in pod color are primarily due to differences in anthocyanin accumulation, which may also be linked to flavor and quality differences in the resulting chocolate (Motamayor *et al*. 2013; Li *et al*. 2021). While there is a wide amount of research involving the agricultural, morphological, and metabolic aspects of *T. cacao*, little is known about the regulatory mechanisms controlling the pigmentation differences across genetic groups. Generally, the structural genes involved in this pathway, such as *CHS*, *CHI*, *F3H*, *DFR*, *ANS*, and *UGT*, are regulated by a transcription factor complex made up of genes encoding proteins belonging to the MYB, bHLH, and WD40 families (Gonzalez *et al*. 2008; Lloyd *et al*. 2017). This MBW complex-based regulation of anthocyanin biosynthesis is well conserved across plants (Choudhary *et al*. 2026). In *T. cacao*, previous co-expression analyses have shown the association between anthocyanin and proanthocyanidin biosynthesis genes (*DFR*, *ANS*, *LAR*, *ANR*, and *UGT*) to the MYB-bHLH-WD40 (MBW) complex (Motamayor *et al*. 2013; Gallego *et al*. 2021).

With the continuing improvement in sequencing technologies, it has been possible to sequence the whole genome of the variety Criollo (Argout *et al*. 2017) and subvariety Matina (Motamayor *et al*. 2013) that belongs to the Amelonado genetic group derived from the Forastero variety. Other projects have investigated additional *T. cacao* cultivars (Nousias *et al*. 2024; Kulesza *et al*. 2024).

Here, we provide genome sequences of two *Theobroma cacao* plants and use the annotated gene sets to investigate flavonoid biosynthesis. Structural genes and transcription factors involved in the flavonol and anthocyanin biosynthesis pathway in *T. cacao* were identified as part of this work.

## Materials & Methods

### Plant Sample and DNA Extraction

The *Theobroma cacao* trees, XX-0-BRAUN-7843067 and SD-0-BONN-19718, used in this study have been cultivated under greenhouse conditions at the Botanical Gardens of TU Braunschweig and the University of Bonn for decades (**Figure 2**). The origin of both trees is unknown. Fresh and healthy leaves were harvested on the same day DNA extraction was performed to ensure high DNA quantity and quality. The DNA extraction of the SD-0-BONN-19718 tree was performed with a previously described CTAB-based protocol (Siadjeu *et al*. 2020; de Oliveira *et al*. 2026). For the leaves obtained from the tree at the Botanical Garden Braunschweig, the DNA extraction was performed with some modifications as described below. Fresh leaf material was subjected to the removal of the midribs and veins, leaving behind only foliar tissue used for extracting DNA. This was done to reduce the polyphenol and polysaccharide content in the sample. About 0.9 g of the foliar tissue was ground into a fine powder using liquid nitrogen.

**Figure 2.**
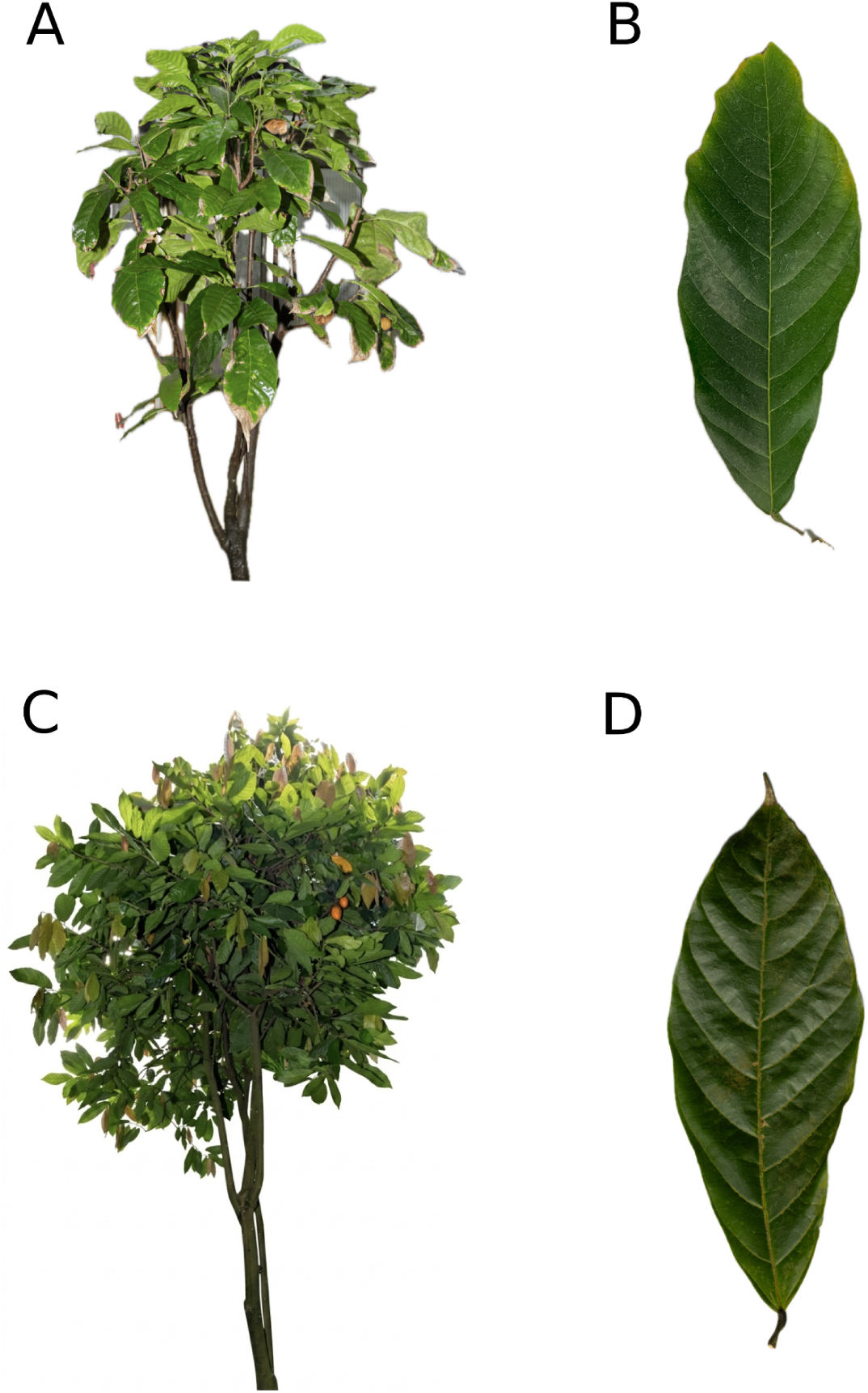
(A) *Theobroma cacao* plant in the Botanical Garden Braunschweig. (B) Adult green leaf from *Theobroma cacao* plant in the Botanical Garden Braunschweig. (C) *Theobroma cacao* plant in the University of Bonn Botanic Gardens. (D) Adult green leaf from *Theobroma cacao* plant in the University of Bonn Botanic Gardens. Photo credits: Jakob Horz.

The sample was vortexed briefly before being introduced into the water bath. The sample was then centrifuged at 11,000 x g at 20 °C for 30 min. After the CTAB2 buffer step, the supernatant was discarded, and a sorbitol wash was performed following the instructions of a previously developed protocol (Souza *et al*. 2012). The sorbitol washing step was repeated only once. After the ethanol wash, the supernatant was again discarded, and the sample was allowed to dry for 5 minutes. Any excess droplets were removed by gently pipetting out and discarding. The sediment was then dissolved in 100 µl CTAB-TE (10 mM Tris, pH 8.0, 0.1 mM EDTA, 7.5 µg/ml of DNAse-free RNase A) and was left to incubate overnight at room temperature. DNA quality was then determined with NanoDrop measurement, agarose gel electrophoresis, and Qubit measurement. Finally, fragments shorter than 25 kb were depleted using the Short Read Eliminator Kit (Pacific Biosciences).

### Nanopore Sequencing

For each library, 1 μg of high molecular weight DNA was used. The DNA was prepared according to the SQK-LSK109 and SQK-LSK114 protocols (ONT), respectively (Additional file 1). Sequencing was conducted using MinIONs with R9.4.1 and R10.4.1 flow cells, respectively. Between sequencing runs, a wash step was performed with the EXP-WSH004 following ONT’s procedures to recover blocked nanopores. For data generated for the plants from the Botanical Garden in Braunschweig, Dorado v6.4.6+ae70e8f (ONT) was used for basecalling with the super high accuracy model (dna_r9.4.1_450bps_sup.cfg and dna_r10.4.1_e8.2_400bps_5khz_sup.cfg). For data generated from the plant material sampled from the Bonn University Botanic Gardens, the basecalling was performed using dorado v1.0.2 and v1.1.1 with the HAC model available for the current versions.

### Genome Sequence Assembly

Genome sequence assemblies were generated using three long-read assemblers: Hifiasm v0.25.0-r726 (Cheng *et al*. 2021), NextDenovo v2.5.2 (Hu *et al*. 2024) and Shasta v0.14.0 (Shafin *et al*. 2020). An estimated genome size of ∼430 Mb was used for all assemblies. For Hifiasm, assemblies were generated using default parameters optimized for ONT reads. Assembly parameters for NextDenovo v2.5.2 with the correct_option included a read cutoff set to 1 kbp, an estimated genome size of 430 Mbp, sort options set to -m 20g -t 15, minimap2_options_raw set to -t 8, pa_correction of 3, and correction_options set to -p 15. The assemble_option was set to minimap2_options_cns - -t 8 and nextgraph_options = -a 1. The assembly parameters for Shasta were based on a pre-built Nanopore-Oct2021 configuration file supplied within Shasta v0.14.0. This configuration uses a minimum read length of 10 kbp and other default parameters defined within the configuration preset, including a k-mer size of 50 with the probability set to 0.05, enrichmentThreshold to 100, and distanceThreshold to 1000. MinHash parameters included version set to 0, m set to 4, hashFraction set to 0.01, minHashIterationCount set to 50, alignmentCandidatesPerRead set to 20, and both minBucketSize and maxBucketSize set to 0.

Completeness of the assembly was assessed using BUSCO v6.1.0 (Simão *et al*. 2015; Manni *et al*. 2021) in the genome mode with eudicotyledons_odb12 (Zdobnov *et al*. 2021; Tegenfeldt *et al*. 2025) as the reference data set. General assembly statistics were calculated with contig_stats3.py, applying a minimum contig length of 10000 bp to exclude shorter contigs. Names of sequences were optimized with clean_genomic_fasta.py to avoid incompatibilities in downstream analysis (https://github.com/bpucker/GenomeAssembly) (de Oliveira *et al*. 2026).

### Comparative genomics

The genome sequences assembled in this study were compared with the previously published genome sequences of the cacao varieties *Criollo* (Argout *et al*. 2017) and *Matina* (Motamayor *et al*. 2013). Transposable elements in all three genomes were annotated using Extensive *de-novo* Annotator (EDTA) v2.2.1 (Ou *et al*. 2026) with consistent parameters: --anno 1 --sensitive 1 --evaluate 1. To exclude gene-related sequences, the coding sequences of the respective genome sequences were supplied using the --cds parameter. NUCMER from the MUMmer v4.0.0rc1 package (Marçais *et al*. 2018) was used to identify the syntenic regions among the genome sequences. RagTag v2.1.0 (Alonge *et al*. 2022) was used to reorder contigs in the assembled genome sequence based on the most contiguous genome sequence.

Tandem repeats in the genome sequences were identified using Tandem Repeats Finder (TRF) v4.09.1 (Benson 1999) using parameters ‘Match=2, Mismatch=5, Delta=7, PM=80, PI=10, Minscore=5 and MaxPeriod=2000’. Telomere repeat units were explored using the Telomere Identification toolkit (TIDK) (Brown *et al*. 2025)for the plant canonical telomeric unit (TTTAGGG/CCCTAAA). A contig/scaffold was classified as T2T if at least 100 occurrences of this repeat were detected at both ends. A Circos plot was generated using Circos v0.69-8 (Krzywinski *et al*. 2009), following previously described methods (Hakim *et al*. 2025).

### Phylogenetic reconstruction

To investigate the evolutionary history of our sequenced cacao plants, genome assemblies of eight *Theobroma cacao* accessions were retrieved from the NCBI database (**Table 1**). Along with the two genome assemblies generated in this study, orthologous gene sets were identified using the BUSCO-based phylogenomics pipeline ArboPhyl v1.1 (Verschuren 2025). BUSCO v5.5 (Manni *et al*. 2021) was run using genome mode with the embryophyta_odb10 lineage dataset. For each genome sequence, BUSCO genes classified as single-copy genes were extracted, only those that shared a threshold of at least 80% sequence similarity across all the genome sequences were retained for downstream analysis. The pipeline extracted the sequences separately for each orthologous group. Sequences of a group were aligned using the alignment tool MAFFT v7.525 (Katoh & Standley 2013) with default parameters. Poorly aligned or ambiguous regions were trimmed using trimAl v1.5 (Capella-Gutiérrez *et al*. 2009), and later concatenated into a supermatrix containing 1443 BUSCO partitions.

**Table 1.**
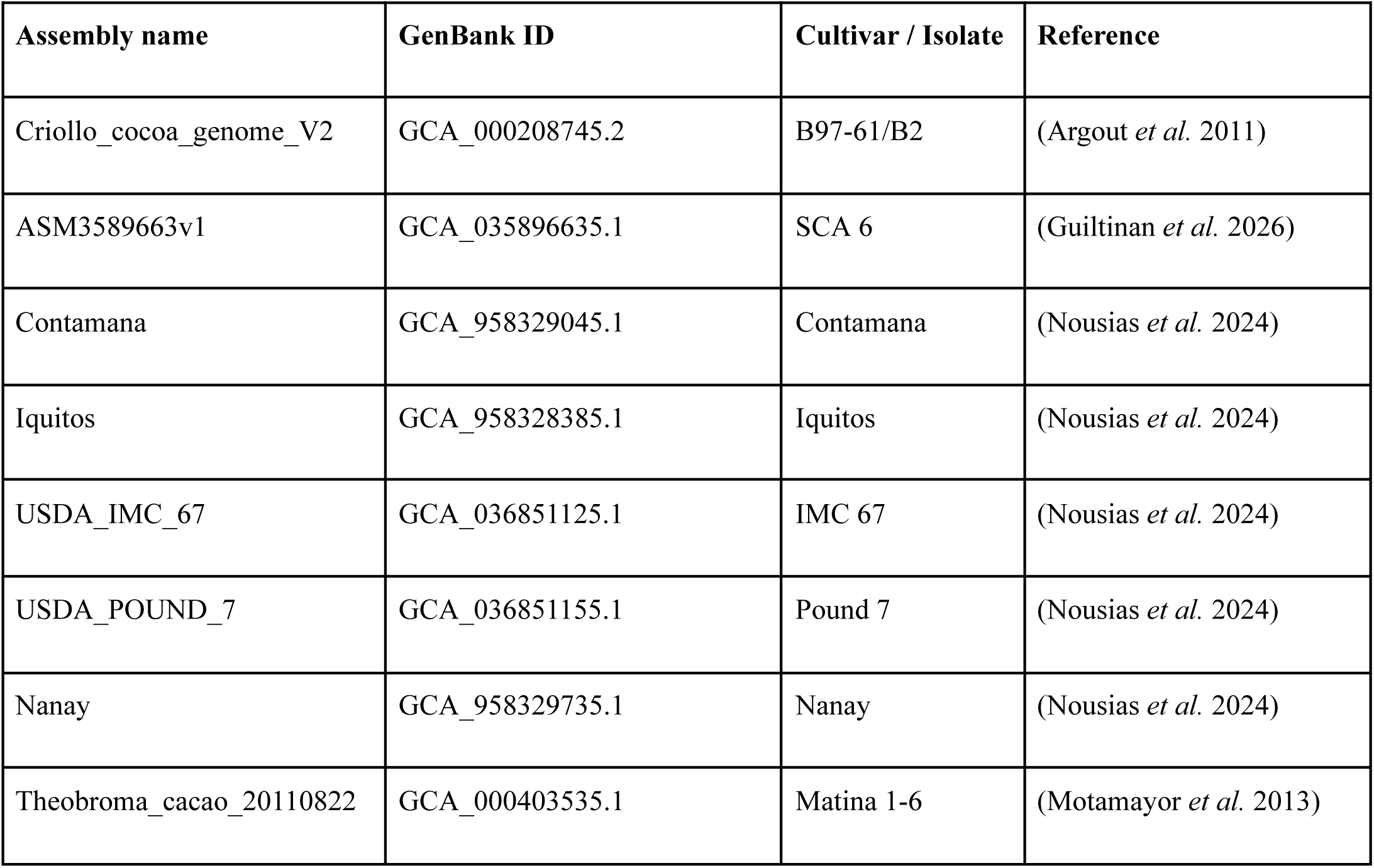
Genome assemblies of *Theobroma cacao* used in the maximum-likelihood phylogenetic reconstruction.

Maximum-likelihood phylogenetic inference was performed using IQ-TREE v3.1.2 (Wong *et al*. 2026). ModelFinder (Kalyaanamoorthy *et al*. 2017) was used to determine the best suitable substitution model for each BUSCO gene using partitions of the alignment (Additional file 2), and phylogenetic reconstruction was conducted using all these individual partitions. Branch support was assessed using 1000 ultrafast bootstrap replicates (Hoang *et al*. 2018).

Additionally, a coalescent species tree was inferred for comparison. For this purpose, individual gene trees were estimated for each BUSCO alignment using IQ-TREE v3.1.2 (Wong *et al*. 2026) under the same partition-specific substitution models mentioned above. The resulting gene trees were then concatenated and used as input for ASTRAL-III v5.7.1 (Zhang *et al*. 2018).

### Structural Annotation

For the prediction of protein-coding genes, GeMoMa v1.9 (Keilwagen *et al*. 2019) was run with the following hints: GCA_035896635, GCA_036851095 (Nousias *et al*. 2024), GCA_036851125 (Nousias *et al*. 2024), GCA_036851155 (Nousias *et al*. 2024), GCF_000208745, and GCF_000403535, and the parameters pc=true pgr=true p=true o=true GeMoMa.c=0.4 GeMoMa.Score=ReAlign Extractor.r=true GAF.f=“start==’M’ and stop==’*’ and (isNaN(score) or score/aa>=’0.75’)” AnnotationFinalizer.r=NO. In addition, a mapping of all available RNA-seq datasets were provided for the generation of additional hints (Additional file 3). The RNA-seq reads were retrieved from NCBI and aligned with STAR v2.7.11b (Dobin *et al*. 2013; Dobin & Gingeras 2015) using default settings. The resulting BAM file was sorted with samtools. The resulting annotation was filtered with the following settings: f=“start==’M’ and stop==’*’ and (isNaN(tie) or tie>0) and tpc>0 and aa>50” atf=“tie>0 and tpc>0”. The annotation results were compared with respect to completeness as measured with BUSCO v6.1.0 in protein mode using eudicotyledons_odb12 lineage (Simão *et al*. 2015; Tegenfeldt *et al*. 2025).

### Functional Annotation and Identification of Flavonoid Biosynthesis Candidate Genes

Functional annotation was performed to identify genes important to the flavonoid biosynthesis pathway. Knowledge-based Identification of Pathway Enzymes (KIPEs) v3.2.6 (Pucker *et al*. 2020; Rempel *et al*. 2023) was applied to identify the structural genes present in the flavonoid biosynthesis pathway based on the FlavonoidBioSynBaits_v3.3. To achieve a detailed annotation of flavonoid UDP-glycosyltransferases (UGTs), polypeptide sequences were collected from GenBank (accession numbers: GCA_000208745.2, GCA_036851125.1, GCA_036851095.1, GCA_036851155.1) and Phytozome (accession number: MD16G1266400). BLASTp was used to identify initial candidates in each *T. cacao* cultivar, with a cutoff of percent identity ≥ 50. MAFFT v7.453 (Katoh & Standley 2013) was used to align the sequences of initial candidates and bait sequences in KIPEs. IQ-TREE v3.1.2 (Wong *et al*. 2026) was used to infer a phylogeny for various glycosyltransferases. iTOL (Letunic & Bork 2021, 2024) was used to visualize and annotate the tree. To further characterize the expression patterns of the identified UGT candidates, transcript abundance data of the varieties Criollo and Matina were filtered to retain candidate genes and visualized as heatmaps (Additional file 4).

Transcript abundance data were generated using various python scripts, all of which are publicly available on GitHub. RNA-seq data were fetched from the Sequence Read Archive (NCBI) using the SRA toolkit (SRA Toolkit Release 3.1.1 (Katz *et al*. 2022)). Transcript abundance was quantified using kallisto 0.51.0 (Bray *et al*. 2016) and count tables were generated using an iterative SRA mapper (Iterative_SRA_mapper.py)(Borchert 2024). The merging, filtration, and co-expression analysis of candidate genes were executed using the co-expression Python script CoExp.py (Pucker & Iorizzo 2023). Heatmaps were generated using a customized Python script with the Seaborn package (Waskom 2021).

MYB_annotator v1.0.1 (Pucker 2022) and bHLH_annotator v1.04 (Thoben & Pucker 2023) were utilized to identify the flavonoid biosynthesis-regulating R2R3-MYB and bHLH genes, respectively. The MYB_annotator.py was run with the AthRefMYBs.txt to perform an orthology detection against *A. thaliana*. The bHLH annotator was run on the pbb-tools web server with default parameters. Initial TTG1 candidates were identified based on sequence similarity to a collection of TTG1 proteins (Pucker *et al*. 2020) via BLASTp (Altschul *et al*. 1990). These candidates were validated through the construction of global alignments with characterized TTG1 and other WD40 sequences via MAFFT. A phylogeny was inferred by FastTree v2.1.10 (Price *et al*. 2010) and inspected via iTOL (Letunic & Bork 2024).

A general functional annotation of all predicted polypeptide sequences was performed with the customized Python script construct_anno.py (Pucker & Iorizzo 2023; de Oliveira *et al*. 2026) based on sequence similarity to characterized *Arabidopsis thaliana* genes as described in TAIR10/Araport11 (Lamesch *et al*. 2012; Cheng *et al*. 2017).

### Structural variation analysis

Structural variants (SVs) were identified in different *T. cacao* accessions by mapping their respective sequencing reads against the Criollo reference genome sequence (GCA_000208745.2) using minimap2 v2.30-r1287. The resulting SAM files were converted to BAM format, sorted, and indexed using SAMtools v1.19.2 (Danecek *et al*. 2021). SVs were detected using Sniffles2 v2.2.1 (Smolka *et al*. 2024) with default parameters. The resulting VCF files containing the identified insertions, deletions, and inversions were visualized using the Integrative Genomics Viewer (IGV) v.2.15.5 (Robinson *et al*. 2011). To assess the potential functional impact of those identified structural variants, candidate genes involved in flavonoid biosynthesis were selected based on the annotations generated using KIPEs v3.2.6 (Pucker *et al*. 2020; Rempel *et al*. 2023). The corresponding protein accessions were used to identify the respective gene models in the Criollo reference genome annotation, from which chromosome, start, end, and strand coordinates were retrieved from the GFF3 annotation. Each candidate-gene region was extended by ±10 kb upstream and downstream to capture regulatory regions. BED files were generated from the annotated gene coordinates and used to subset the BAM files for visualization in IGV (Robinson *et al*. 2011). To evaluate the potential effects of coding-region structural variants at a protein level, polypeptide sequences were used for protein structure prediction. Three-dimensional protein models were generated using Colabfold v1.6.2 (Mirdita *et al*. 2022).

### Identification of Transcription Factor Binding Sites (TFBSs) in flavonoid biosynthesis gene promoters

To investigate structural variation affecting the regulation of flavonoid biosynthesis genes among the cacao accessions, the promoter regions corresponding to six genes (*CHS1, FNSII, F3H, FLS1, DFR2, and LAR1*) were extracted from the reference *T. cacao* Criollo genome sequence (GCA_000208745.2), and the newly generated genome sequences. Putative promoter regions were defined as 2 kb upstream of the inferred transcription start site (TSS). TSS positions were inferred from the annotated mRNA coordinates and strand orientation in each corresponding GFF3 annotation file. Genomic coordinates were converted to BED format and sequences were extracted using BEDTools v2.31.0 (Quinlan & Hall 2010) getfasta with the strand-aware (-s) option to ensure all sequences were reported in the transcriptional 5’-3’ orientation. SV coordinates were compared with the 2 kb promoter regions using BEDTools intersect.

Variants overlapping promoter regions were considered candidate TFBS structural variants and were further evaluated based on their position relative to gene orientation and predicted impact on transcriptional regulation. For those candidate SVs overlapping the putative promoter regions in the *T. cacao* accessions studied here, the original sequence was extracted from the Criollo reference genome sequence and screened for known plant transcription factor binding motifs using the MEME Suite tool, FIMO v5.5.9 (Grant *et al*. 2011). Motif occurrences were analyzed using a *T. cacao* transcription factor binding motif dataset from the PlantTFDB v5.0 (Guo *et al*. 2008).

## Results

### Assembly Results

Different assemblies produced as part of this study were compared against each other and against the available genome sequences of previously investigated cultivars (**Table 2**). As mentioned earlier, two *T. cacao* plants were selected as sources for genome sequencing: one from the Botanical Garden in Braunschweig and one from the Botanical Garden in Bonn. The resulting genome assemblies are hereafter referred to as BRAU1 and BONN1, respectively. Three long-read genome assemblers, Hifiasm, NextDenovo, and Shasta were used to independently assemble each genome, resulting in three assemblies for BRAU1 and three assemblies for BONN1, giving a total of six *de novo* genome assemblies. All six assemblies presented very high completeness values, with BUSCO scores ranging from 98.9% to 99.8% of completeness based on eudicot reference genes (Additional File 5). Among these, the BRAU1 and BONN1 Hifiasm assemblies showed the best overall performance with BUSCO completeness scores of 99.1 % and 99.8%, respectively. Additionally, both Hifiasm assemblies presented the greatest contiguity with N50 values of 37.4 Mbp and 45.6 Mbp (**Table 2**). The BONN1 genome assembly represented a near T2T gap-free genome sequence based on the identification of putative centromere-associated loci based on tandem repeat enrichment. 19 out of the 20 telomeres with thousands of copies of telomeric repeat units (TRUs) were identified, suggesting that 9 out of 10 contigs were T2T **(Figure 3)**. Therefore, these assemblies were selected as the representative genome sequences and subjected to functional annotations of the predicted genes. High macrosynteny was observed between the 10 pseudochromosomes of *Matina* and *Criollo* varieties and the top 13 and 10 largest pseudochromosomes of the BRAU1 and BONN1 cacao genome sequences, respectively, which collectively represent 95% and 96% of the respective assembly sizes.

**Figure 3.**
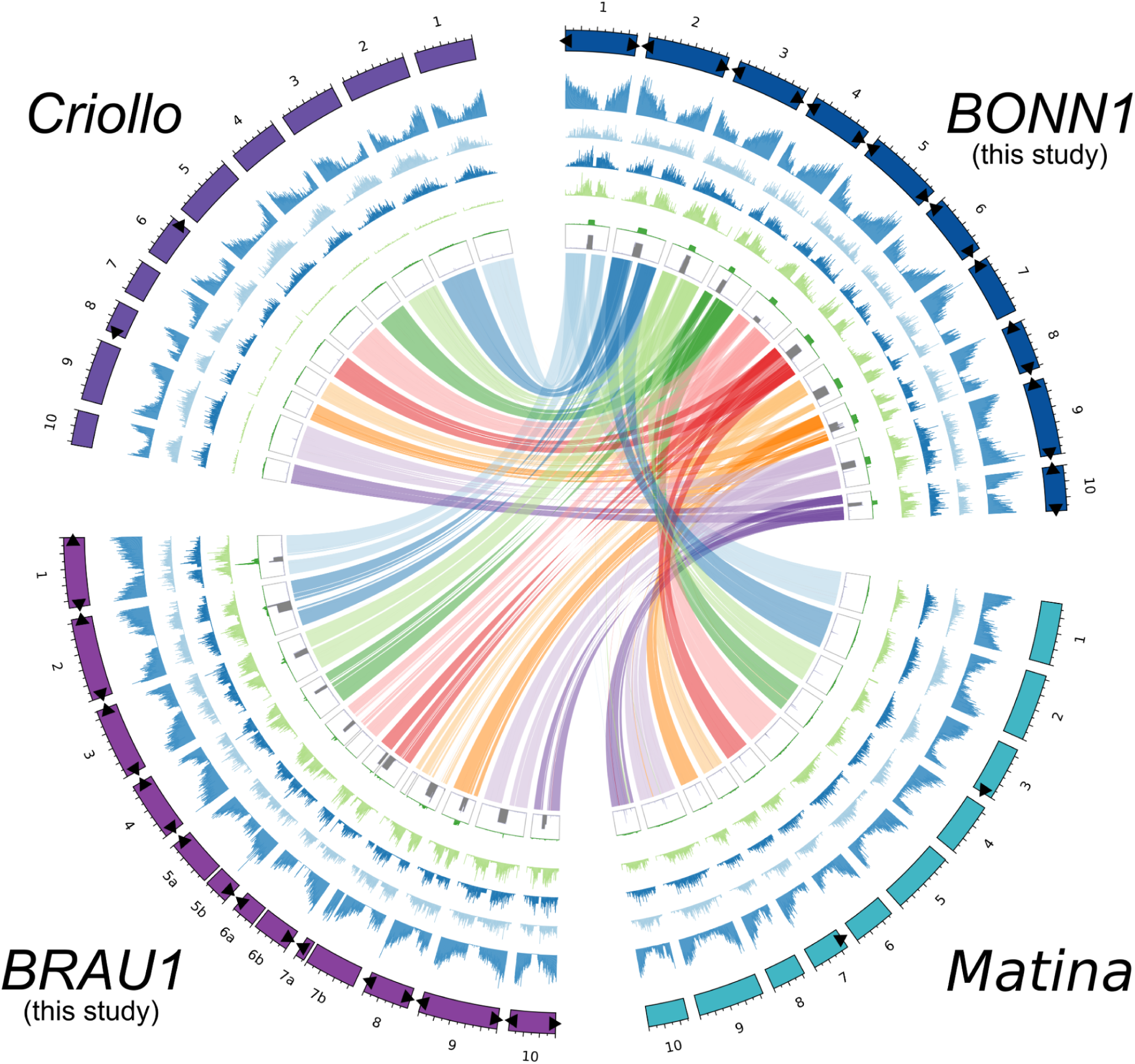
Circos plot comparing the sequenced *Theobroma cacao* genomes in this study, BONN1 and BRAU1 with the *Matina* and *Criollo* (Motamayor *et al*. 2013; Argout *et al*. 2017) varieties. From the outermost to innermost layers: circular representation of top longest contigs/scaffolds (colored ideograms) from from each assembly, densities of protein-coding genes, gypsy-type LTR retrotransposons, copia-type LTR retrotransposons, TIRs, tandem repeat density, and macrosynteny among the three genome sequences with respect to BONN1 genome sequence. All density information is calculated in non-overlapping 100-kbp windows. Black triangles mark identified telomeric sequences on the ends of the sequences.

**Table 2.**
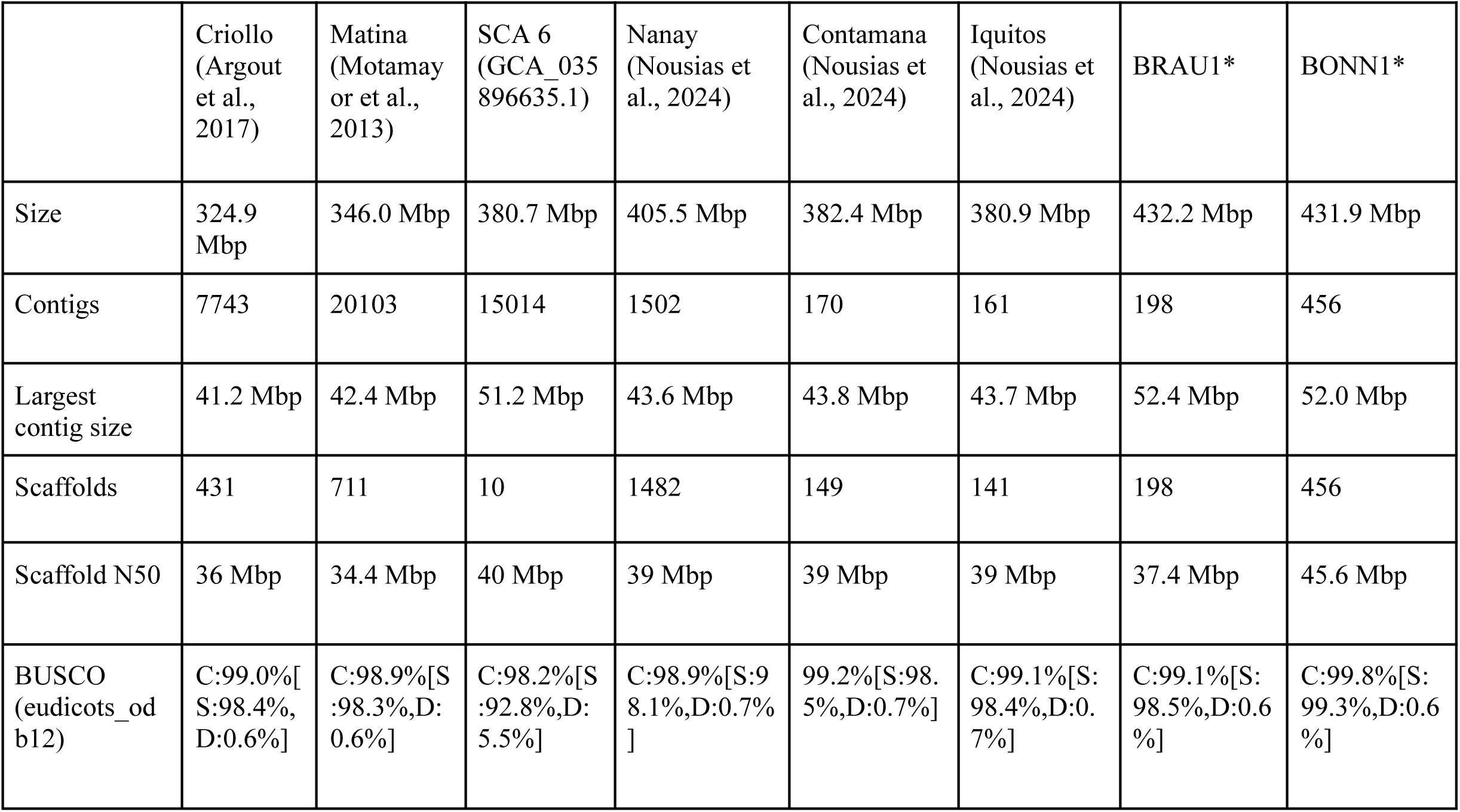
Statistics of different *T. cacao* genome sequences. An asterisk indicates the representative genome sequence produced by Hifiasm in this study.

|  | Criollo<br>(Argout<br>et al.,<br>2017) | Matina<br>(Motamay<br>or et al.,<br>2013) | SCA 6<br>(GCA_035<br>896635.1) | Nanay<br>(Nousias et<br>al., 2024) | Contamana<br>(Nousias et<br>al., 2024) | Iquitos<br>(Nousias et<br>al., 2024) | BRAU1* | BONN1* |
| --- | --- | --- | --- | --- | --- | --- | --- | --- |
| Size | 324.9<br>Mbp | 346.0 Mbp | 380.7 Mbp | 405.5 Mbp | 382.4 Mbp | 380.9 Mbp | 432.2 Mbp | 431.9 Mbp |
| Contigs | 7743 | 20103 | 15014 | 1502 | 170 | 161 | 198 | 456 |
| Largest<br>contig size | 41.2 Mbp | 42.4 Mbp | 51.2 Mbp | 43.6 Mbp | 43.8 Mbp | 43.7 Mbp | 52.4 Mbp | 52.0 Mbp |
| Scaffolds | 431 | 711 | 10 | 1482 | 149 | 141 | 198 | 456 |
| Scaffold N50 | 36 Mbp | 34.4 Mbp | 40 Mbp | 39 Mbp | 39 Mbp | 39 Mbp | 37.4 Mbp | 45.6 Mbp |
| BUSCO<br>(eudicots_od<br>b12) | C:99.0%[<br>S:98.4%,<br>D:0.6%] | C:98.9%[S<br>:98.3%,D:<br>0.6%] | C:98.2%[S<br>:92.8%,D:<br>5.5%] | C:98.9%[S:9<br>8.1%,D:0.7%<br>] | 99.2%[S:98.<br>5%,D:0.7%] | C:99.1%[S:<br>98.4%,D:0.<br>7%] | C:99.1%[S:<br>98.5%,D:0.6<br>%] | C:99.8%[S:<br>99.3%,D:0.6<br>%] |

### Annotation of the *Theobroma cacao* genome sequence

The predicted protein-coding genes in the *Theobroma cacao* BRAU1 and BONN1 genome assemblies showed BUSCO completeness of 99.1% for both annotations. In the BRAU1 annotation, a total of 43,582 genes were predicted, with an average protein length of 423 amino acids (aa) and an average coding sequence (CDS) length of 1,270 bp. The BONN1 annotation contained a total of 45,753 predicted genes, with average protein and CDS lengths of 375 aa and 1,124 bp, respectively.

The BRAU1 and BONN1 assemblies contained similar proportions of transposable elements (TEs), accounting for 52.6% and 52.4 % of the respective genome assemblies. However, the composition of the TE landscape differed between the two accessions. Retrotransposons accounted for 81% of all repetitive sequences in BRAU1, compared with 57% in BONN1, with long terminal repeat-retrotransposons (LTR-RTs) accounting for most of this difference. In contrast, Mutator TIRs accounted for approximately 17% of the BONN1 genome but only ∼5% of the BRAU1 genome (**Figure 3)** Both newly sequenced genomes contained a higher proportion of repetitive sequences than the previously reported Matina (44%) and Criollo (46%) genomes. This difference may partly reflect the larger assembled genome sizes and the improved recovery of complex repetitive regions enabled by long-read sequencing. Retrotransposons represented the predominant TE class in both previously reported genomes, accounting for 77.2% and 72% of repetitive sequences in Matina and Criollo, respectively (Additional file 6).

### Phylogeny of *Theobroma cacao* cultivars inferred from orthologous genes

To determine the phylogenetic placement of BRAU1 and BONN1 cacao plants, concatenated and coalescent species trees were reconstructed based on single-copy orthologous genes shared among the newly assembled genome sequences and those previously published *Theobroma cacao* genome assemblies (**Figure 4**). Overall, the two approaches presented strong bootstrap support for most of the major clades.

**Figure 4.**
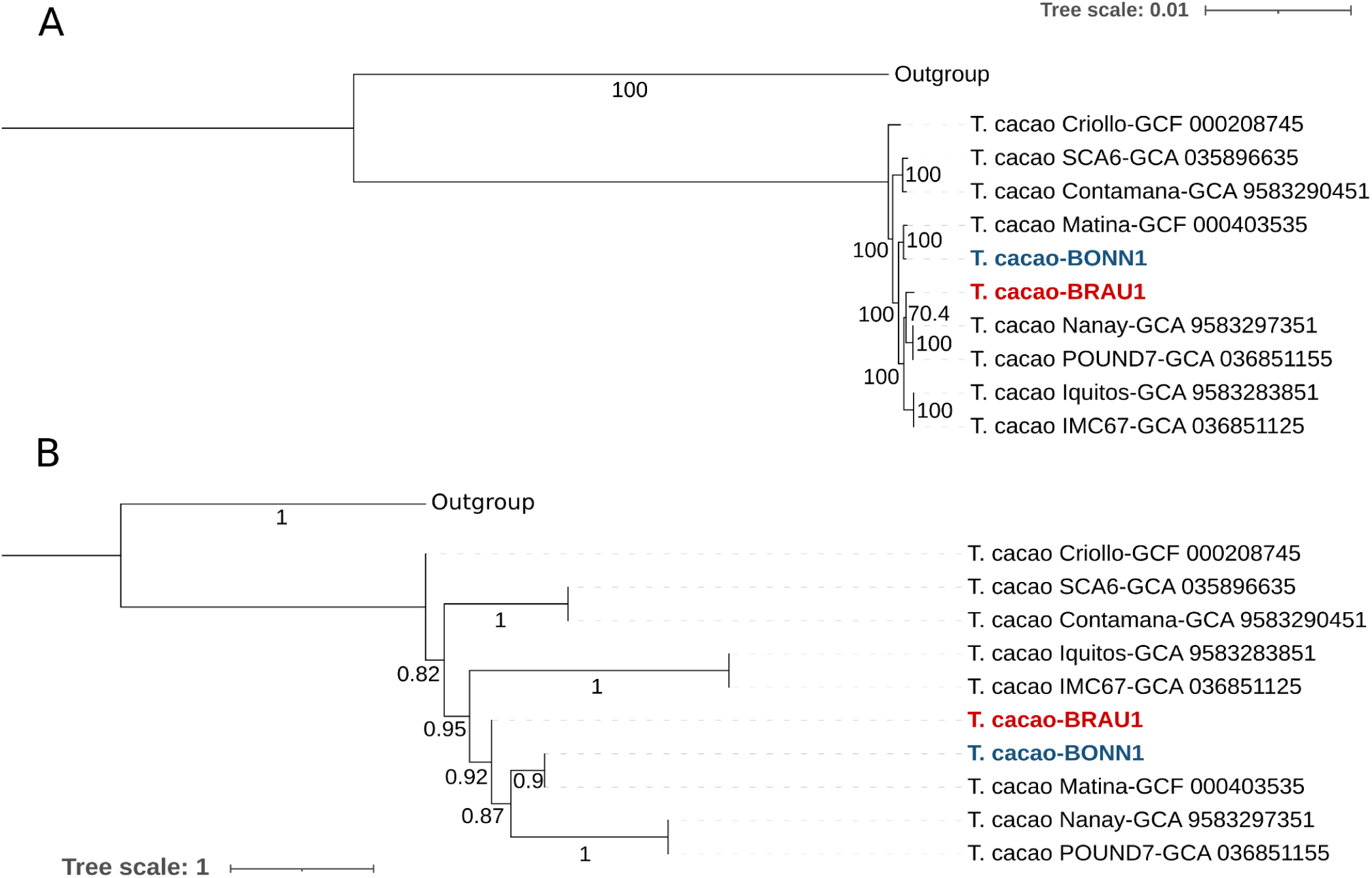
Phylogenetic relationships among the newly assembled *Theobroma cacao* genomes BRAU1 and BONN1 and previously published *T. cacao* genome assemblies. (A) Concatenated maximum-likelihood phylogenetic tree reconstructed from single-copy orthologous BUSCO genes. Node values represent bootstrap support from 1000 replicates with values ranging from 0 to 100. (B) ASTRAL coalescent species tree reconstructed from the same set of single-copy orthologous BUSCO genes. Numbers above the nodes are the bootstrap values (ranging from 0-1) of maximum likelihood (ML) analysis based on 1000 replicates. In both trees, BRAU1 and BONN1 are highlighted in red and blue, respectively. Both trees were rooted using *Gossypium* species as the outgroup.

The newly assembled BRAU1 and BONN1 genome sequences were consistently clustered within the lineage containing the Matina, Nanay, and Pound7 accessions, indicating a closer evolutionary relationship with these cultivars. Notably, in both trees Criollo occupies a basal position relative to the remaining *T. cacao* accessions, separated from the clade containing SCA6, Contamana, Iquitos, IMC67, and the newly assembled genome sequences. This topology indicates that Criollo represents the earliest diverging lineage among the sampled taxa.

### Candidate Genes of the Flavonoid Biosynthesis Pathway

Structural genes of the general phenylpropanoid and core flavonoid biosynthesis pathways, as well as key transcriptional regulators were identified within the gene set predicted for BRAU1 and BONN1 (**Figure 5**). These structural genes included *PAL, C4H, 4CL, CHS, CHI, FNSII, F3’H, F3’5’H, F3H, FLS, DFR, ANS, LAR, ANR, arGST,* and *UGT*. While some central genes such as *CHI, F3H, ANS, arGST, DFR,* and *ANR* are present as single-copy genes, there are several genes showing evidence of duplication, including *PAL, C4H, 4CL, CHS, FLS,* and *LAR*.

**Figure 5.**
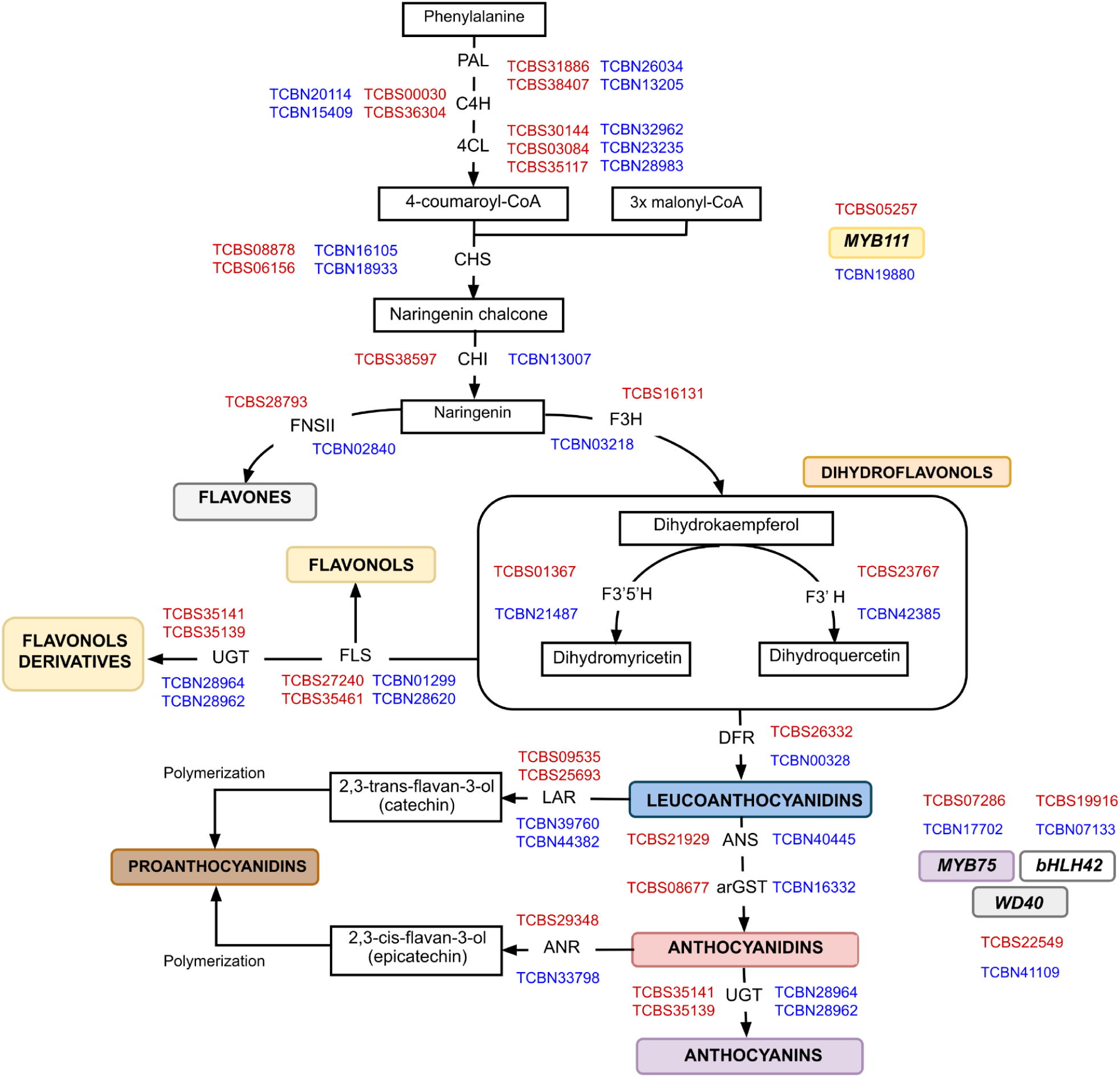
Schematic illustration of the flavonoid biosynthesis with cacao candidate genes of BRAU1 (red) and BONN1 (blue). The displayed structural genes include phenylalanine ammonia-lyase (*PAL*), cinnamic acid 4-hydroxylase (*C4H*), 4-coumarate-CoA ligase (*4CL*), chalcone synthase (*CHS)*, chalcone isomerase (*CHI)*, flavanone 3-hydroxylase (F3H), flavonol synthase (*FLS)*, dihydroflavonol 3-reductase (*DFR)*, anthocyanidin synthase/leucocyanidin dioxygenase (*ANS/LDOX)*, leucoanthocyanidin reductase (*LAR)*, anthocyanidin reductase (*ANR)*, anthocyanin-related glutathione S-transferase (*arGST)*, flavonoid 3’-hydroxylase (*F3’H)*, flavonoid 3’,5’-hydroxylase (*F3’5’H)*, flavone synthase II (F*NSII)*, and flavonoid-dependent glycosyltransferase *(UGT)*. Additional modification enzymes are listed in Additional file 4. In addition, the transcription factors MYB111 (flavonol regulator), MYB75/PROMOTER OF ANTHOCYANIN PRODUCTION 1, bHLH/TRANSPARENT TESTA 8, WD40/TRANSPARENT TESTA GLABRA 1 (anthocyanin regulator) are shown. An asterisk next to a gene identifier indicates that the candidate lacks an expected amino acid residue in a functionally important position.

### Comparative analysis of flavonoid gene models reveals structural divergence among cacao genomes

To investigate the contribution of structural variations (SVs) to genomic diversity among cacao accessions, an SV calling analysis was performed using the Criollo genome sequence as reference. This analysis revealed specific insertions and deletions affecting six key genes involved in flavonoid biosynthesis, including *CHS, FNSII, F3H, FLS, DFR,* and *LAR* (Additional file 7). The identified structural variants were distributed within the specific targeted genes or in their surrounding genomic regions and varied considerably in size, ranging from small insertions and deletions to large genomic deletions exceeding 6 kb.

Among the analyzed loci, the largest structural variations were detected in *DFR2* and *LAR1* (Additional file 7), where deletions of 1026 bp and 6690 bp, respectively, were identified. These larger SVs represent strong candidates for influencing flavonoid pathway variation due to their potential effects on gene structure and regulation. In *DFR2,* the identified deletion results in the loss of two annotated exons and the promoter region, in which TF binding sites (TFBSs) have been predicted (**Figure 6A**). The gene prediction process resulted in protein sequences in BRAU1 (294 aa) and BONN1 (222 aa) that were considerably shorter than the one predicted for the Criollo reference (367 aa). This difference in the number of encoded amino acids indicates loss of a large portion of the protein (**Figure 6B**). To further characterize the *DFR* copies, protein sequence alignments were performed for DFR1 from Criollo, BRAU1, BONN1, and Matina together with the identified duplicated DFR2 from Criollo, BRAU1, and BONN1 (**Figure 6C**). DFR1 showed a greater sequence conservation across the analyzed cacao genome sequences, whereas DFR2 displayed higher sequence divergence from DFR1. Interestingly, the amino acid corresponding to position 193 in *T. cacao* DFR1, which is known to be a residue associated with DFR substrate specificity, differed between the two DFR copies. DFR1 contained an asparagine (N) residue at this position, while DFR2 contained an alanine (A) residue. Based on a previously established classification (Choudhary & Pucker 2024), these residues correspond to DFR_N_ and DFR_A_ types, respectively. Therefore, the two DFR copies show differences not only in their genomic structure but also in their predicted protein sequences, including a residue associated with substrate preference.

**Figure 6.**
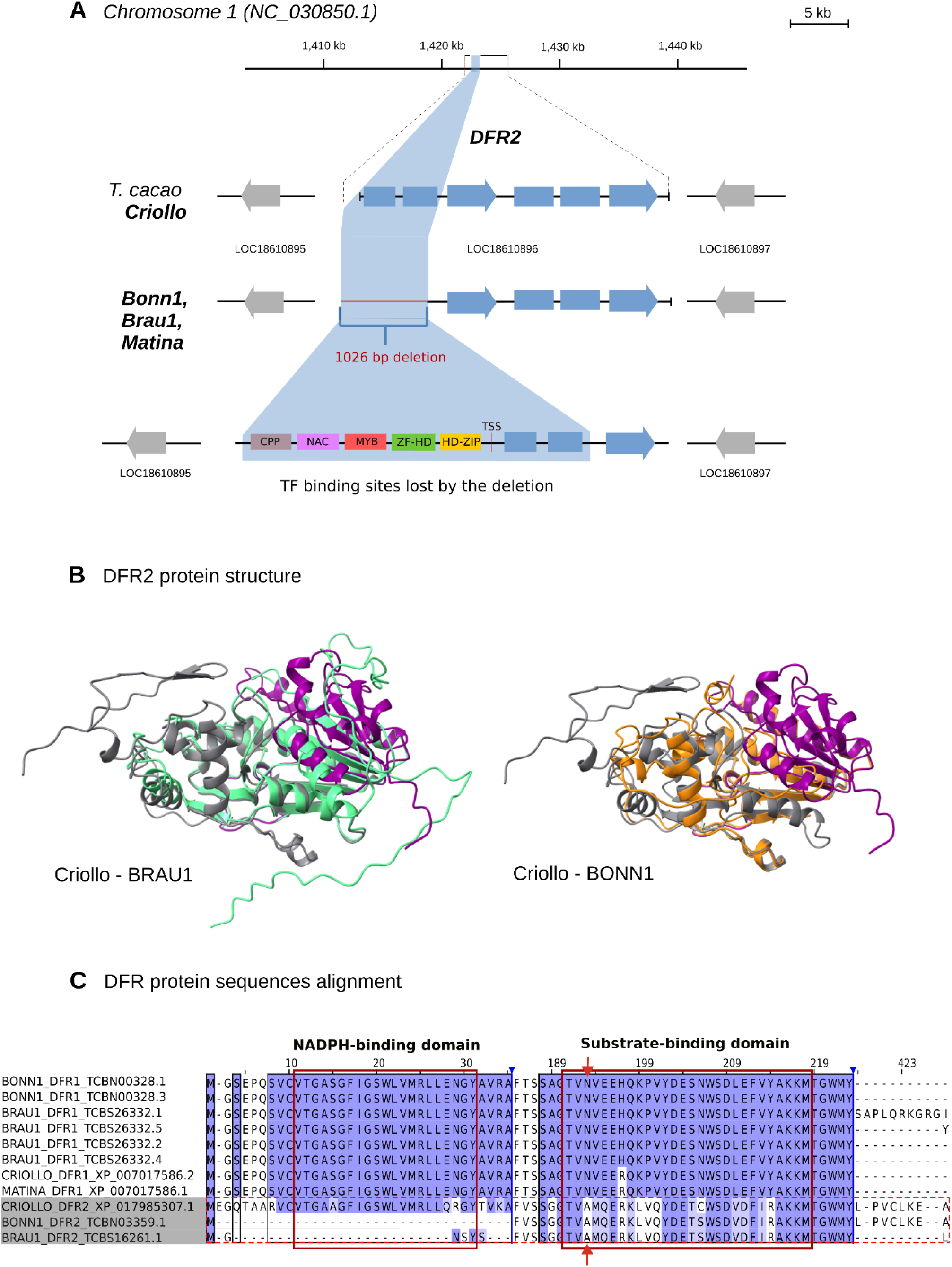
Structural variation and predicted protein structure of DFR2 alleles. (A) Schematic representation of the *DFR2* genomic locus showing the structure of the gene in the reference Criollo and the location of a 1026 bp deletion among the BRAU1, BONN1 and Matina accessions. Predicted Transcription Factor Binding Sites (TFBSs) present in Criollo and potentially lost by the deletion are indicated at the bottom. Exons are represented by blue boxes and introns by connecting lines. The arrows indicate the transcription orientation and flanking genes. (B) Predicted structural models of the DFR2 protein in Criollo (gray), BRAU1 (green), and BONN1 (orange). The location of the deletion in the protein is highlighted in purple. (C) Multiple sequence alignment of DFR protein sequences. Sequence similarity is indicated by purple shading according to the BLOSUM62 similarity score, with darker shades representing higher similarity. The NADPH-binding region and the substrate-binding region are highlighted in red boxes. The residue at the third position of the substrate-binding region, associated with dihydroflavonol substrate recognition, is indicated with red arrows. DFR1 contains an N residue at this position, corresponding to the DFR_N_ type, whereas DFR2 contains an A residue, corresponding to the DFR_A_ type, following a previously established classification (Choudhary & Pucker 2024). The alignment was generated using MAFFT v7.505 and visualized using IGV v.2.15.5.

## Discussion

### Contiguous genome sequences

The genome sequences enabled a comprehensive identification of protein-coding genes. The genome sequences reported in this study are characterized by substantially fewer contigs than previously reported genome sequences of other cultivars (Motamayor *et al*. 2013; Argout *et al*. 2017; Nousias *et al*. 2024; Kulesza *et al*. 2024). The Matina 1-6 genome sequence listed in **Table 1** was assembled by combining data from NGS technologies and BAC sequences. Contigs were sorted and oriented to construct pseudochromosomes for microbial resistance analysis. The Criollo B97-61/B2 genome sequence listed in **Table 1** is also based on multiple genomic libraries. These short read sequencing approaches differ from the long read genome sequencing approach selected in this study because the DNA was sequenced exclusively using ONT. This may explain why the genome sequence produced by this study has higher contiguity i.e., fewer contigs while the total assembly size is larger. This observation is in line with previous studies observing the superiority of long read sequencing over short read sequencing (Marks *et al*. 2021; Pucker *et al*. 2022). The number of protein-coding genes in the *T. cacao* genome sequence matches the average number of genes previously reported in other plant genome sequences (Pucker & Brockington 2018). The high percentage of recovered BUSCO reference genes also supports a high quality of the provided gene set. In summary, it appears that the genome sequence assembled in this study and the corresponding annotation are well equipped for genome-scale analyses, such as identifying the genetic basis of pathogen resistances, yield, taste, or other traits.

### Gene duplication in the flavonoid biosynthesis

On the basis of a comprehensive structural annotation, genes involved in flavonoid biosynthesis were identified and subjected to further investigation, including expression analysis. The discovery of all genes required for the core flavonoid biosynthesis steps aligns with previous reports about the detection of flavonoids in *T. cacao* (Ramiro *et al*. 2005; Gallego *et al*. 2021; Li *et al*. 2021; Hermund *et al*. 2024).

Previous studies have identified genes of the flavonoid biosynthesis in different *T. cacao* varieties (Gallego *et al*. 2021; Li *et al*. 2021). While central genes like *ANS* and *ANR* appear with a single copy, there are several cases of gene duplication events, e.g., involving *FLS*, *DFR*, and *LAR*. Gene duplication is a common phenomenon underlying the complexity of the specialized plant metabolism (Ober 2005; Panchy *et al*. 2016; Lichman *et al*. 2020; Pucker & Iorizzo 2023). Following a gene duplication, the two emerging copies can follow different evolutionary trajectories, often resulting in sub- or neofunctionalization.

### Structural variation in *T. cacao* genomes

The comparative analysis performed in this study revealed that all six investigated flavonoid biosynthesis genes annotated in BRAU1 and BONN1 exhibited structural variation relative to the Criollo reference genome, although the frequency, size, and distribution of these variants differed considerably among loci (Additional file 7). Structural variations (SVs), including insertions, deletions, inversions, duplications, and translocations larger than 50 bp, have been widely studied as a major source of genetic diversity within plant genomes (Saxena *et al*. 2014; Alonge *et al*. 2020). While less studied than single-nucleotide polymorphism (SNPs), SVs have received increasing attention lately due to improvements in sequencing technologies, particularly the expansion of long-read sequencing (Yuan *et al*. 2021). Additionally, since SVs include larger modified fragments of DNA, their effects have a greater potential to affect the structure and expression of the gene by altering their coding and regulatory regions, respectively. This makes SVs important drivers of phenotypic diversity, plasticity, and domestication across multiple crop species such as tomato (Alonge *et al*. 2020), rapeseed (Zhang *et al*. 2024), and cacao (Hämälä *et al*. 2021). Pangenome studies of SVs in *T. cacao* have demonstrated that insertions and deletions contribute substantially to the genetic differentiation among cultivated and wild cacao populations, affecting both coding and non-coding regions and contributing to variation in important agronomic traits involved in pathogen resistance (Hämälä *et al*. 2021). Cacao presents a complex domestication history with a wide geographical distribution of its germplasm that has given rise to highly divergent genetic groups (Bekele *et al*. 2006; Zhang & Motilal 2016; Nieves-Orduña *et al*. 2023). Therefore, as previously demonstrated, structural variants play an important role in shaping phenotypic diversity, including differences in fruit pigmentation (You *et al*. 2026), yield (Song *et al*. 2023), disease resistance (Jobson & Roberts 2022), and metabolite composition (Li *et al*. 2023).

### Intraspecific *DFR* variation in *T. cacao*

Among all the flavonoid biosynthesis genes analyzed in this study, *DFR2* presented the strongest evidence of a potentially biologically relevant SV. The 1026 bp deletion detected in BRAU1, BONN1, and Matina 1-6 was associated with alterations in gene structure, coinciding with the loss of two annotated exons relative to the Criollo reference genome sequence. This resulted in a considerably different coding sequence that prevented the locus from being annotated as DFR in those accessions while indicating the presence of a duplicated DFR locus in Criollo. As a key gene in anthocyanin and proanthocyanidin biosynthesis, the *DFR2* deletion represents a strong candidate identified in this study with potential functional diversity of the flavonoid biosynthesis pathway among the analyzed cacao accessions. The crucial role of DFR in plant pigmentation has been demonstrated across diverse angiosperm lineages, mutations affecting *DFR* expression or coding sequence have been directly linked to altered anthocyanin accumulation in *Ipomoea* (Tanaka *et al*. 2004), *Petunia* (Liu *et al*. 2026), *Arabidopsis thaliana* (Shirley *et al*. 1992), and *Solanum* species (Wang *et al*. 2022). Therefore, recent studies have identified *DFR* as the structural gene most frequently associated with pigmentation differences (Marin-Recinos & Pucker 2024). DFR is often considered the initial enzymatic step in the synthesis of anthocyanins and proanthocyanidins and participates in substrate competition with FLS in the formation of flavonols (Choudhary *et al*. 2026). Consequently, a blockade or interruption of DFR will direct the substrate influx towards FLS and the synthesis of flavonols, which can reduce pigmentation (Luo *et al*. 2016). Most importantly, the deletion affected not only part of the coding sequence, but removed the promoter region of the *DFR2* gene, causing the loss of predicted TF binding sites. Anthocyanin biosynthesis-activating MYBs, as part of the MBW complex, play a crucial role in the regulation of genes in the flavonoid biosynthesis (Gonzalez *et al*. 2008; Marin-Recinos & Pucker 2024; Choudhary *et al*. 2026). The cis-regulatory elements considered vital for the binding of the MYB protein and consequent activation of transcription have been lost through the deletion in DFR2. Therefore, this SV may affect both the integrity of the encoded protein, if formed, and the transcriptional regulation of *DFR2*. We speculate that expression of this DFR locus has been lost due to the deletion. However, a second *DFR* copy remains intact in both genome sequences, BRAU1 and BONN1, suggesting that the deletion of *DFR2* may not result in the complete loss of the DFR function in the plant. The presence of these duplicated *DFR* loci could be the result of an evolutionary duplication event followed by subfunctionalization. This premise is supported by evidence of divergence at a residue associated with substrate specificity. Choudhary and Pucker classified DFR proteins based on the amino acid present at position 133 in the *Arabidopsis thaliana* DFR, distinguishing DFR_N_, DFR_D_, and DFR_A_ types. DFR_N_ is characterized by an asparagine (N) residue and DFR_A_ by an alanine (A) at this position (Choudhary & Pucker 2024). Alignment of DFR sequences from Criollo, Matina, BRAU1, and BONN1 revealed that DFR1 contains an N residue at the equivalent position (193 in *T. cacao* DFR), which corresponds to the DFR_N_ type, whereas the duplicated DFR2 contains A193, corresponding to the DFR_A_ type. The authors further proposed that DFR_D_ originated from a duplication of DFR_N_, with this duplication being observed in the Fabales and Malvales clades (Choudhary & Pucker 2024). As *T. cacao* belongs to the Malvales order, the presence of two divergent *DFR* copies is consistent with a broader history of DFR duplication and diversification within this taxon. Interestingly, our identification of a DFR_A_-type *DFR2* in *T. cacao* also extends the occurrence of this DFR type beyond the Rosales and Alismatales clades (Choudhary & Pucker 2024).

Additionally, our phylogenetic analysis places Criollo as a basal lineage relative to the other *T. cacao* accessions. Because Criollo retains an intact DFR_A_-type *DFR2*, whereas a large deletion was identified at the corresponding locus of that gene in BRAU1, BONN1 and Matina, it raises the possibility that the DFR_A_-type DFR2 represents a retained copy of an ancestral duplicated *DFR* locus that was subsequently lost or degraded in other cacao lineages. In this context, the deletion of *DFR2* in BRAU1, BONN1 and Matina may represent the loss of a functionally differentiated DFR copy, while the remaining DFR_N_-type DFR1 may retain the core DFR function. However, comparative expression analyses would be required to determine whether the two cacao *DFR* copies have indeed undergone subfunctionalization.

## Conclusions

In this study, two high-contiguity *Theobroma cacao* genomes were sequenced and assembled, BRAU1 and BONN1, both including complete structural and functional annotations that enabled genome-scale analysis of flavonoid biosynthesis. Comparative analysis identified the major structural and regulatory genes of the pathway. Structural variation analysis further revealed insertions and deletions in six flavonoid biosynthesis genes, with a particularly important 1026 bp deletion at the *DFR2* locus in BRAU1, BONN1, and Matina accessions. This deletion was identified to remove predicted promoter elements and two exons in the gene with potential effects on *DFR2* regulation and protein function in the plant. However, an intact *DFR1* copy was also annotated. When comparing the peptide sequences of DFR1 and DFR2, a divergence at a residue associated with substrate preference was identified. These findings suggest that the duplicated copies may have followed distinct evolutionary trajectories contributing to flavonoid diversity in cacao.

## Declarations

## Data availability statement

All data sets underlying this study are publicly available. Sequencing data have been deposited at the European Nucleotide Archive (PRJEB65101 and PRJEB97086, Additional File 1). The assembled genome sequence, corresponding annotation, and analyzed gene expression data have been published via bonndata (https://doi.org/10.60507/FK2/YQDFT4). Customized Python scripts for the analyses in this study are available through GitHub (https://github.com/bpucker/CacaoGenomics).

## Author contributions

MFMR and BP designed the project and supervised the work. KL optimized the DNA extraction protocol for *Theobroma cacao* and performed high molecular DNA extraction. KL, KW, RF, JAVSdO, and CMD performed nanopore sequencing. MFMR, SW, KL, BA, NC, and BP performed data analysis. MFMR, SW, KL, and BP wrote the initial draft of the manuscript. All authors revised the manuscript and agreed to its submission. Order of authorship is based on the time point of joining the project.

## Acknowledgements

This work was supported by the de.NBI Cloud within the German Network for Bioinformatics Infrastructure (de.NBI) and ELIXIR-DE (Forschungszentrum Jülich and W-de.NBI-001, W-de.NBI-004, W-de.NBI-008, W-de.NBI-010, W-de.NBI-013, W-de.NBI-014, W-de.NBI-016, W-de.NBI-022). We thank all members of the research group Plant Biotechnology and Bioinformatics for discussion and support. We are grateful to Jakob Horz for kindly providing the plant photographs included in this manuscript. We are grateful for the excellent support provided by the team of the University of Bonn Botanic Gardens.

## Additional information

**Additional File 1:** List of all ONT sequencing data sets belonging to the *Theobroma cacao* plants sequenced in this study. All data sets are available from the European Nucleotide Archive (ENA).

**Additional File 2**: Best-fitting substitution models identified by ModelFinder for each BUSCO partition used in the phylogenetic analysis.

**Additional File 3**: RNA-seq data sets that served as hints for the GeMoMa annotation of protein-coding genes. All data sets are available from the Sequence Read Archive (SRA).

**Additional File 4**: Glycosyltransferase genes involved in flavonol and anthocyanin modification and their expression patterns in the *Theobroma cacao* varieties Matina and Criollo.

**Additional File 5**: Comparison of assembly statistics for previously published *Theobroma cacao* genome assemblies and six de novo genome assemblies generated in this study using Hifiasm, NextDenovo, and Shasta.

**Additional file 6:** Summary of repeat content identified in the genome sequence of previously reported Matina and Criollo varieties as well as in BONN1 and BRAU1 sequenced and assembled in this study.

**Additional file 7:** Structural variations identified in the flavonoid biosynthesis genes *CHS1, FNSII, F3H, FLS1, LAR1,* and *DFR2* in the *Theobroma cacao* genomes BRAU1 and BONN1 relative to the Criollo reference genome.

