## Additional file 4 for "*Theobroma cacao* genome sequence reveals insights into flavonoid biosynthesis"

### Glycosyltransferases involved in flavonol and anthocyanin modification

Additionally to the expression analysis, a hierarchical cluster analysis was performed to illustrate the relationships among genes based on their gene expression patterns, as result genes were separated into distinct clusters. Among them, the expression patterns of the glycosyltransferase genes, *F3GT* and *A5GT* in the two *Theobroma cacao* varieties, Matina and Criollo, formed clear clusters with other flavonoid genes, providing insights into their roles in the modification of anthocyanins and flavonols (**Additional file 4**). In Matina, each one of the three *F3GT* copies cluster independently, *arGST* clustered along with $F3GT_1$, $FLS_2$ with $F3GT_2$ and $F3GT_3$ formed a bigger cluster with $FLS_1$, and the anthocyanin transcription factor *MYB75* and the flavonol transcription factor *MYB111.* Moreover, the expression of *LAR*, *DFR*, and *ANR* is dominant in certain samples, suggesting a focus on proanthocyanidin biosynthesis (**Additional file 4A**). The moderate expression of *F3GT* and *A5GT* alongside *FLS* indicates that the glycosylation process in Matina may primarily target flavonols rather than anthocyanins. This aligns with Matina's apparent preference for the proanthocyanidin pathway.

In Criollo, *F3GT* and *A5GT* are strongly co-expressed and cluster tightly with genes like *FLS, F3'H, LAR* and anthocyanin transcription factor *MYB75* and flavonol transcription factor *MYB111.* The independent co-clustering of $F3GT_{1,2}$ with $FLS_1$, $MYB111_{1,2,3}$ and $F3'H_2$, and $A5GT_{1,2}$ with $MYB111_4$, $F3'H_1$, and $FLS_2$, indicates that these glycosyltransferases are functionally linked to flavonol biosynthesis. The observed high expression of genes like *ANR, DFR,* $A5GT_1$, $F3'H_1$, and $MYB111_4$, suggests that multiple branches of the flavonoid biosynthesis pathway are active in Criollo (**Additional file 4B**). Similar to Matina, the high expression of *DFR*, *ANS*, and *ANR* shows a dominance in the proanthocyanidin over anthocyanin and flavonol production, however the clustering in expression patterns supports the glycosylation of mainly flavonols over anthocyanins.

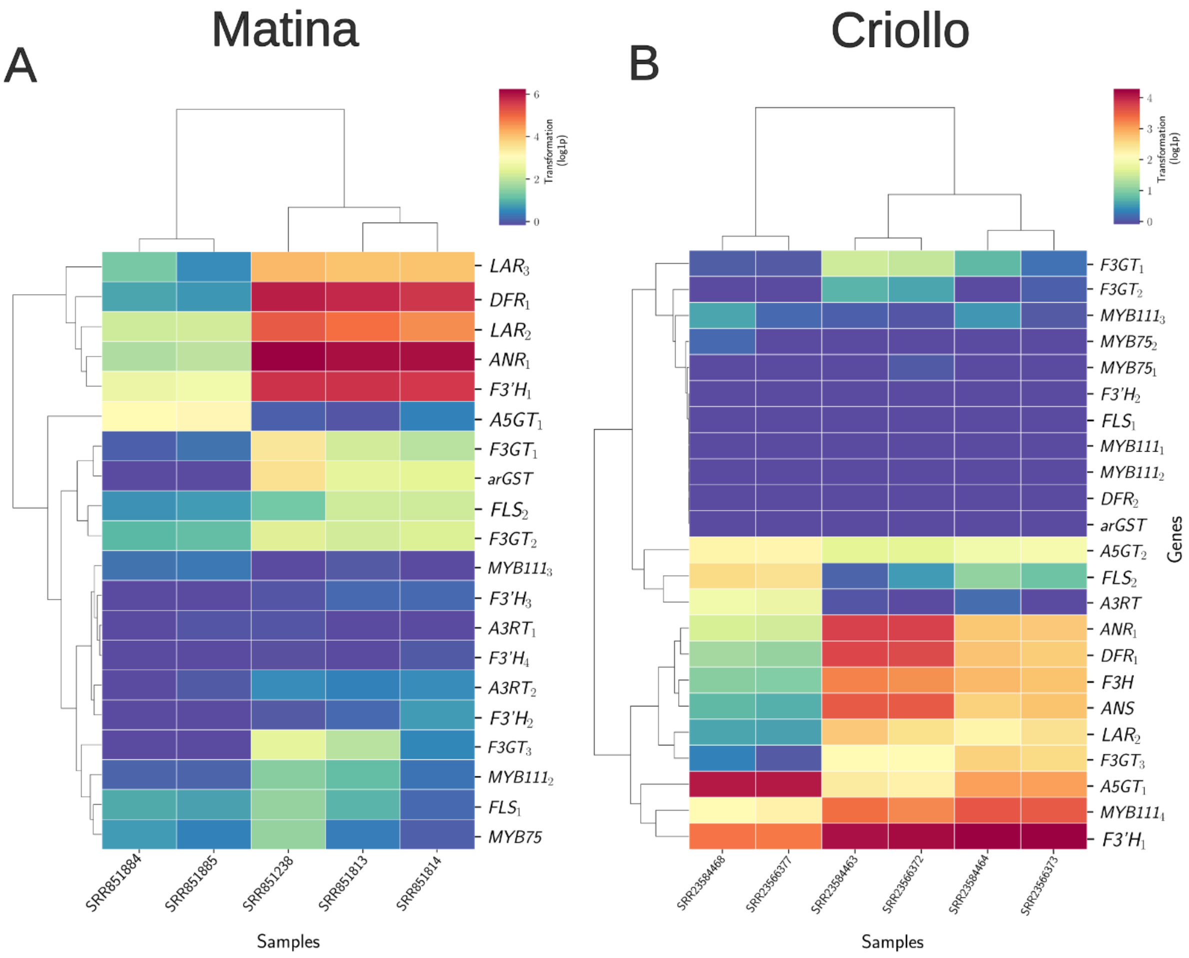


**Additional file 4.** Hierarchical clustering of genes based on different expression levels. Variety Matina with leaf samples (A) and variety Criollo with leaf samples (B). Displayed genes are chalcone synthase (*CHS)*, chalcone isomerase (*CHI)*, flavanone 3-hydroxylase (*F3H*), flavonoid 3'-hydroxylase (*F3'H)*, flavonoid 3',5'-hydroxylase (*F3'5'H)*, dihydroflavonol 4-reductase (*DFR)*, anthocyanidin synthase/leucocyanidin dioxygenase (*ANS/LDOX)*, anthocyanin-related glutathione S-transferase (*arGST)*, flavonol synthase (*FLS)*, leucoanthocyanidin reductase (*LAR)*, anthocyanidin reductase (*ANR)*, and flavonoid glucosyltransferase (*F3GT*), *MYB111* (FlavonolMYB), *MYB75/PROMOTER OF ANTHOCYANIN PRODUCTION 1*.
