## Additional file 5 for "*Theobroma cacao* genome sequence reveals insights into flavonoid biosynthesis"

Comparative assembly metrics of *Theobroma cacao* genome assemblies. Previously published reference genome sequences are compared with the six *de novo* assemblies generated in this study (highlighted in red) using three long-read assemblers (Hifiasm, NextDenovo, and Shasta).

| | Criollo (Argout et al., 2017) | Matina (Motamayor et al., 2013) | SCA 6 (GCA_035896635.1) | Nanay (Nousias et al., 2024) | Contamana (Nousias et al., 2024) | Iquitos (Nousias et al., 2024) | **Hifiasm (This study - BRAU1)** | **Hifiasm This study - BONN1)** | **NextDenovo (This study - TUBRAU)** | **NextDenovo (This study - UBonn)** | **Shasta (This study - TUBRAU)** | **Shasta (This study - UBonn)** |
|---|---|---|---|---|---|---|---|---|---|---|---|---|
| **Size** | 324.9 Mbp | 346.0 Mbp | 380.7 Mbp | 405.5 Mbp | 382.4 Mbp | 380.9 Mbp | 432.2 Mbp | 431.9 Mbp | 372 Mbp | 349 Mbp | 444 Mbp | 405.9 Mbp |
| **Contigs** | 7743 | 20103 | 15014 | 1502 | 170 | 161 | 198 | 456 | 115 | 39 | 575 | 183 |
| **Scaffolds** | 431 | 711 | 10 | 1482 | 149 | 141 | 198 | 456 | 115 | 39 | 575 | 183 |
| **Scaffold N50** | 36 Mbp | 34.4 Mbp | 40 Mbp | 39 Mbp | 39 Mbp | 39 Mbp | 37.4 Mbp | 45.6 Mbp | 12.8 Mbp | 20 Mbp | 8.3 Mbp | 12.5Mbp |
| **BUSCO (eudicots_odb12)** | C:99.0% [S:98.4%, D:0.6%] | C:98.9% [S:98.3%, D:0.6%] | C:98.2% [S:92.8%, D:5.5%] | C:98.9% [S:98.1%,D:0.7%] | 99.2% [S:98.5%, D:0.7%] | C:99.1% [S:98.4%, D:0.7%] | C:99.1% [S:98.5%, D:0.6%] | C:99.8% [S:99.3%, D:0.6%] | C:98.9% [S: 97.8%, D:1.2%] | C:99.8% [S: 99.1%, D:0.7%] | C:98.9% [S:97.8%,D:1.2%] | C:99.8% [S:91.2%, D:8.6%] |
