## Additional file 7 for "*Theobroma cacao* genome sequence reveals insights into flavonoid biosynthesis"

### Structural variation at six flavonoid biosynthesis genes in *Theobroma cacao* genome sequences.

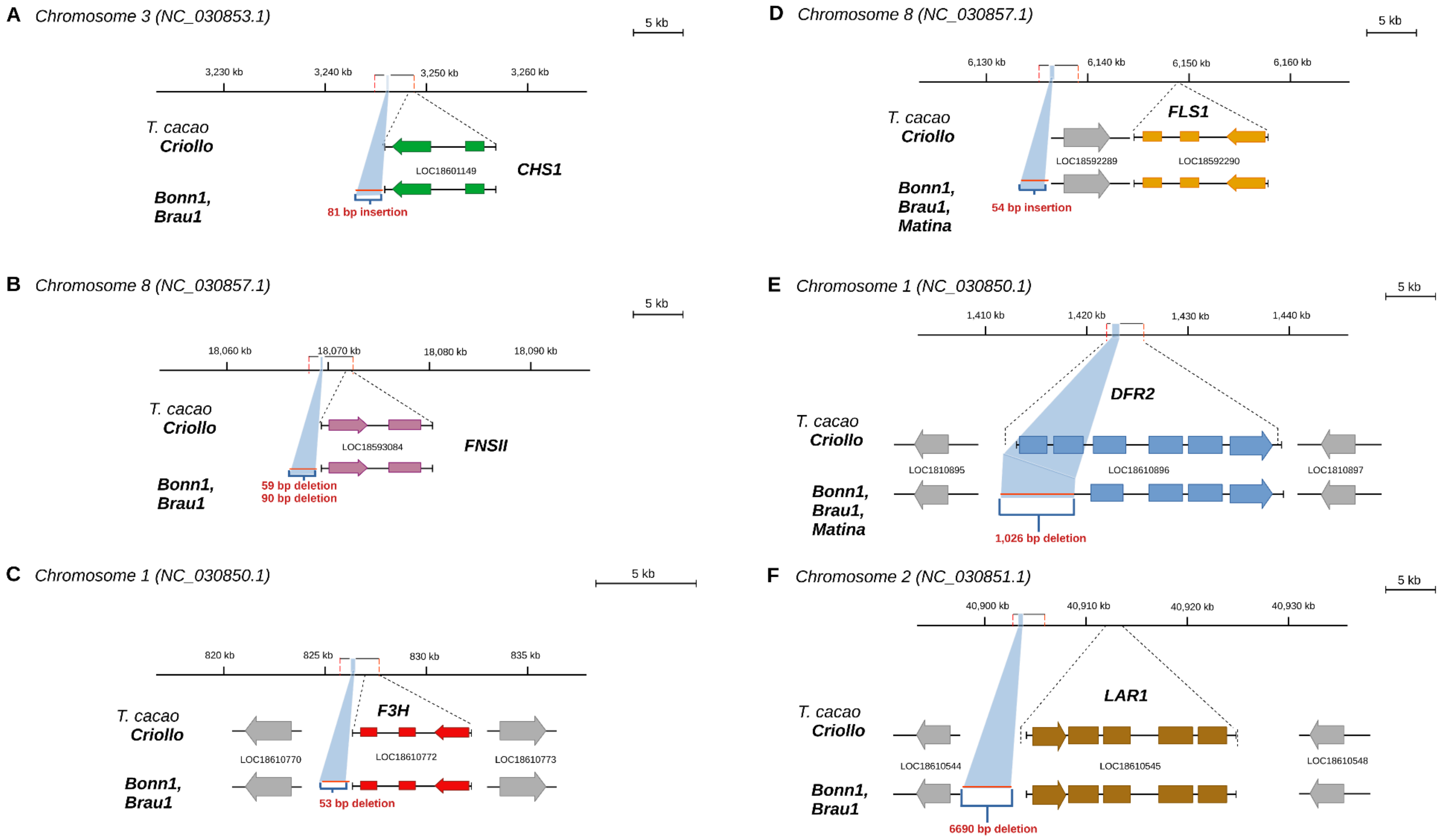


Schematic representation of the genomic regions surrounding: (A) *CHS1*, (B) *FNSII*, (C) *F3H*, (D) *FLS1,* (E) *LAR1* and (F) *DFR2* in the Criollo reference genome sequence and the BONN1 and BRAU1 genome sequences. Gene models and structural variants are shown relative to the Criollo reference coordinates. Insertions and deletions identified between the genome sequences are indicated by their corresponding sizes.
